# Capture and visualization of live *Mycobacterium tuberculosis* bacilli from tuberculosis bioaerosols

**DOI:** 10.1101/2019.12.23.887729

**Authors:** Ryan Dinkele, Sophia Gessner, Anastasia S. Koch, Carl Morrow, Melitta Gqada, Mireille Kamariza, Carolyn R. Bertozzi, Brian Smith, Courtney McLoud, Andrew Kamholz, Wayne Bryden, Charles Call, Valerie Mizrahi, Robin Wood, Digby F. Warner

**Affiliations:** SAMRC/NHLS/UCT Molecular Mycobacteriology Research Unit & DST/NRF Centre of Excellence for Biomedical TB Research, Department of Pathology, Faculty of Health Sciences, University of Cape Town, South Africa; Institute of Infectious Disease and Molecular Medicine, Faculty of Health Sciences, University of Cape Town, South Africa; Desmond Tutu HIV Centre, University of Cape Town, South Africa; Department of Biology, Stanford University, CA 94305, USA; Department of Chemistry, Stanford University, CA 94305, USA; Howard Hughes Medical Institute, Stanford University, CA 94305, USA; Edge Embossing, USA; Zeteo Tech, USA; Wellcome Centre for Infectious Disease Research in Africa, University of Cape Town, South Africa

## Abstract

The tuberculosis (TB) pandemic demands urgent interventions such as those designed to interrupt *Mycobacterium tuberculosis* (*Mtb*) transmission, a challenge exacerbated by our poor understanding of the events enabling successful transfer of infectious bacilli between hosts. To address this problem, we developed the Respiratory Aerosol Sampling Chamber (RASC), a personal clean-room equipped with high-efficiency filtration and sampling technologies that allow biosafe capture and isolation of particulate matter – including *Mtb* bacilli – released by patients during natural breathing and (non-induced) cough. Here, we demonstrate the use of DMN-trehalose labelling to detect and quantify live *Mtb* bacilli among complex bioaerosol samples arrayed in a bespoke nanowell device following capture in the RASC. A pilot study identified *Mtb* in more than 85 % of known TB patients, improving significantly on previous work which has relied on animal infection and cough sampling to estimate transmission events. Moreover, intra-patient comparisons of bioaerosol and sputum samples indicated that *Mtb* aerosols likely derive from a compartment other than sputum. These results support the utility of the RASC platform for research aimed at interrupting *Mtb* transmission, including the non-invasive detection of *Mtb*-infected individuals who are predicted to contribute to bacillary spread despite the absence of clinical symptoms.

## Introduction

*Mycobacterium tuberculosis* (*Mtb*) is the causative agent of tuberculosis (TB), the leading infectious killer globally having claimed 1.4 million lives in 2018 (*1*). TB control is almost entirely predicated on the treatment of active disease. However challenges posed by the long duration of standard chemotherapy (*2,3*), delayed and missed diagnoses (*4,5*), and failure to retain patients in treatment programmes, mean the goals of global TB eradication remain aspirational (*6*). The propagation of multi-drug resistant (MDR) disease further imperils this approach; an assessment reinforced by growing evidence of the primary transmission of MDR *Mtb* strains (*7*–*12*) and the knowledge that MDR-TB now accounts for more than one-quarter of all antimicrobial resistance (AMR) cases annually. These factors, together with the increasing appreciation that asymptomatic *Mtb* infections (with or without underlying pathology) might contribute to bacillary dissemination (*13*), have re-focused attention on *Mtb* transmission as a critical, but poorly understood, area for novel interventions in endemic regions (*14*–*16*). It is sobering, for example, that only 1-30 % of new *Mtb* infections can be linked to known TB cases – a disconnect which implies the existence of many unrecognized transmitters in TB endemic communities (*17*–*20*).

An obligate pathogen, *Mtb* must drive successive cycles of transmission, infection, and disease in order to persist within the human population. Therefore, while the interval between host infection and disease can be variable (*21*–*23*) – perhaps involving extended periods of clinical latency in some cases (*24*–*26*)– the biological imperative to effect successful transmission (thus ameliorating the risks of extinction in an individual host) is inexorable and has been proposed to underlie the destructive lung pathology which enables aerosolization and release of *Mtb* bacilli (*21,27,28*). Studying aerosolized *Mtb* is complicated by the low numbers of bacilli produced and the presence of environmental and patient-derived “contaminating” microorganisms and particulate matter (*29*). While molecular methods have enabled consistent and reliable detection of *Mtb* DNA in bioaerosols (*30*–*32*), these approaches cannot distinguish live from dead organisms nor do they allow investigations of the physiological state(s) of the aerosolized bacilli. The method of bacillary capture is also key: approaches based on forced or induced cough might not reflect natural transmission events (*33,34*), whereas face-mask and related aerosol sampling methodologies either make live-cell analyses impossible (*35*) or require downstream *in vitro* propagation (via microbiological culture) (*36*), unavoidably altering the physiological and metabolic state of the captured samples.

In attempting to address these limitations, we developed a platform for the detection and capture of live *Mtb* bacilli from TB patients (*37*). Our Respiratory Aerosol Sampling Chamber (RASC) effectively functions as a personal clean-room, enabling collection and concentration of particulate material released by an individual patient during normal respiratory activity, including from natural (non-induced) cough (*29,37*). Through multiple iterations informed by our preliminary experimental and clinical observations, we have equipped the RASC to address systematically the comprehensive TB transmission research agenda outlined by others (*38*). Key questions posed include: When are viable, infection-competent bacilli released by *Mtb*-infected individuals? How many aerosolized bacilli are produced, and are they single *Mtb* bacilli and/or clumps? Do clinically latent TB cases transmit *Mtb*? What is the impact of anti-TB chemotherapy on production of viable *Mtb* aerosols? Does sputum smear-positivity predict infectiousness? Are sputum bacilli representative of the transmitted bacillary population? Is cough essential for transmission? Are some *Mtb* genotypes (better) adapted to transmission? How long do bacilli remain viable following aerosolization? Are some individuals “super-spreaders” with an increased propensity for *Mtb* bioaerosol production? To date, we have demonstrated the potential for liquid capture of aerosolized *Mtb* in the RASC, eliminating the dependency on solid culture-based techniques for bacillary detection (*29,37*). This represents a key development, since it promises the possibility of detecting, isolating, and manipulating live bacilli for downstream phenotypic and genomic studies of transmission. For that, however, optimized protocols are needed to enable the specific labelling and concentration of low numbers of *Mtb* bacilli from mixed samples, and in a format amenable to detection via live-cell microscopy and, ultimately, experimental analysis.

The solvatochromic dye, 4-*N,N*-dimethylamino-1,8-napthalimide-trehalose (or DMN-tre), was developed as a fluorescent probe for the rapid detection of *Mtb* in sputum samples (*39,40*). Enzymatic incorporation of this trehalose analogue into the hydrophobic mycomembrane by the mycobacterial Antigen-85 complex, as either a trehalose mono- or di-mycolate, enhances DMN fluorescence nearly one thousand-fold, limiting background noise attributable to unincorporated probe, and circumventing the need for multiple wash steps (*39,41*). This is a key attribute of DMN-tre and facilitates its use for the detection of *Mtb* bacilli within sputum in under an hour (*39,41*). Importantly, since active membrane biosynthesis is a pre-requisite for bacterial labelling, DMN-tre possesses the additional advantage of detecting live organisms only.

In this study, we combine the bioaerosol sampling capabilities of the RASC (*37*) with DMN-tre fluorescence detection (*39*) for the visualization and image-based phenotypic characterization of live *Mtb* bacilli in bioaerosol samples obtained from TB patients. We demonstrate that DMN-tre labelling of samples captured and concentrated in liquid enables fluorescence-dependent visualization of organisms in specialized nanowell arrayed devices. Then, in a pilot study, we use this system to detect putative *Mtb* in more than 85 % of confirmed TB patients sampled, discerning three broadly distinct mycobacterial phenotypes associated with aerosolization. In addition, we compared matched bioaerosol and sputum samples to elucidate differences in labelling profiles. Our results establish the capacity for sensitive, non-invasive detection of infectious bioaerosols from TB-active individuals. Importantly, they also provide essential proof-of-concept data in support of current work which aims to extend sampling to asymptomatic, potentially *Mtb*-positive individuals in addressing critical programmatic and scientific questions that must be tackled before interventions to interrupt TB transmission in endemic settings can be considered.

## Results

### Liquid cyclone capture, nanowell arraying and DMN-tre detection of TB bioaerosols

We previously described the capacity to enumerate *Mtb* genomes (via RD9-specific PCR) or *Mtb* colonies (via growth on solid medium loaded into different impactor devices) captured in bioaerosol material produced by tidal breathing and (non-induced) cough during the ∼60 minute isolation of confirmed TB patients in the RASC (*29,37*). Although instructive, both methods (molecular and microbiological) are characterized by limitations which critically undermine their respective utilities for TB transmission research. On the one hand, PCR cannot differentiate live from dead bacilli, thereby rendering any estimate of viable *Mtb* release uncertain. On the other, culture-based methods are notoriously susceptible to contamination, the complications arising from the presence of so-called “viable but non-culturable” or “differentially detectable” organisms (*29,42*), and the temporal – and, consequently, genetic (*43*) and physiological – separation of the (single) transmitted bacillus from the micro-population (∼10^6^ cells) which eventually manifests as a visible *Mtb* colony on a plate. For these reasons, we deemed culture-independent detection, quantification and visualization of live organisms a priority.

Recent innovations in the RASC technology have ensured that approximately 500 L of expired air per subject per hour can be concentrated into ∼5-10 mL sterile phosphate-buffered saline (PBS). This development, which is enabled through the use of a Bertin Coriolis μ Biological Air Sampler, is significant in allowing high sampling volumes while producing a tractable, low-volume liquid output for downstream manipulation and analysis. Owing to its described utility for detecting *Mtb* in sputum samples from TB patients (*39*), DMN-tre was selected as the preferred label for the specific detection of *Mtb* within concentrated bioaerosol samples arrayed on a bespoke nanowell device (**Figs. 1A & 1B**). The superstructure of each device consisted of two rows of eight wells (16 in total) overlaid on an embossed cyclic olefin copolymer (COC) film. The COC film contained the 50 × 50 µm nanowells which were arrayed ∼140 µm apart center-to-center (**Fig. 1C**). Each microwell therefore comprised approximately 1600 nanowells.

**Fig. 1.**
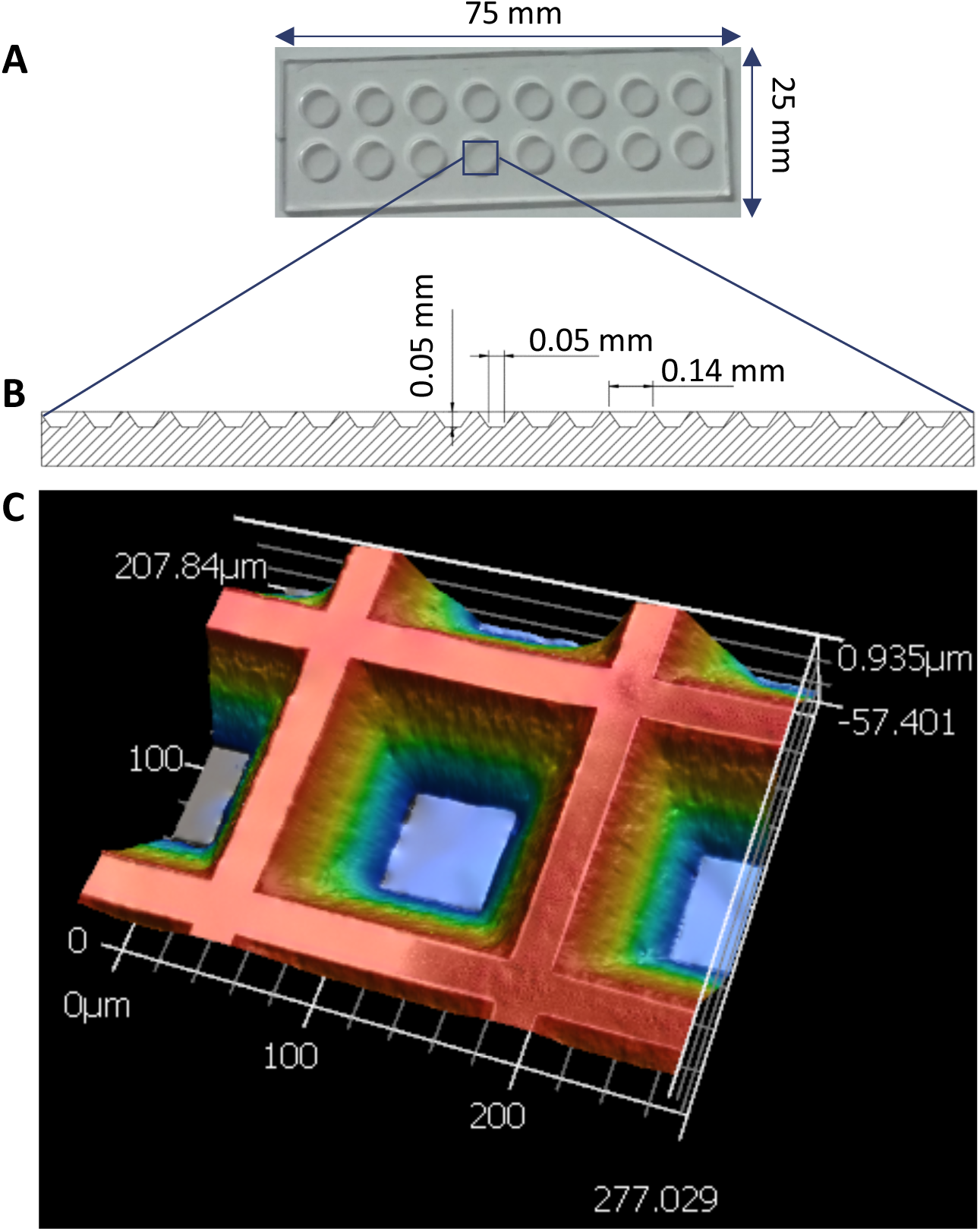
Design and fabrication of nanowell-arrayed microscope slides for the compartmentalization and visualization of TB bioaerosols. (**A**) Photograph of a nanowell slide. (**B**) Schematic of the nanowell device. (**C**) 3D scan of a 207.840 × 277.029 µm section of the slide. Each device (25 × 75 mm) consists of two rows of eight round microwells machined from cast acrylic. The microwells are 6 mm in diameter and 2 mm deep. The nanowell film, which is adhered to the superstructure via UV-curing adhesive, is made from embossed COC film. The nanowells have side-wall angles of 35° and are 50 µm deep. The distance through the bottom of each well to the back of the film is roughly 170 µm, equivalent to a number 1.5 coverslip.

Following incubation in the presence of DMN-tre, the entire concentrated aerosol sample from a single patient was added to a single microwell. Thereafter, brief centrifugation of the device ensured dispersal of the liquid sample across thousands of individual nanowell compartments for fluorescence imaging. Physical separation of samples across the nanowells was considered a potential benefit in ensuring that all viable organic material was isolated in discrete wells, thereby facilitating detection while, potentially, reducing the likelihood that faster-growing non-*Mtb* organisms (“contaminants” for TB diagnostic purposes, but natural components of the patient aerosol microbiomes) would overwhelm the device following overnight DMN-tre labelling.

### Microscopic analyses revealed the complexity of bioaerosol samples

To evaluate this technology, we initially recruited into a pilot study only TB-confirmed patients who had yielded GeneXpert-positive, drug-susceptible samples at presentation. Following routine diagnosis, the patients were transferred to the research site for RASC sampling; ethical approval sanctioned the delayed initiation of standard TB chemotherapy for 2 hours to enable the bioaerosol work. The ∼5-10 mL Bertin liquid-capture sample was concentrated via centrifugation before resuspension in 200 μl standard Middlebrook 7H9 growth medium supplemented with 100 μM DMN-tre for overnight (16 h) incubation at 37°C to allow metabolic incorporation (**Fig. 2**).

**Fig. 2.**
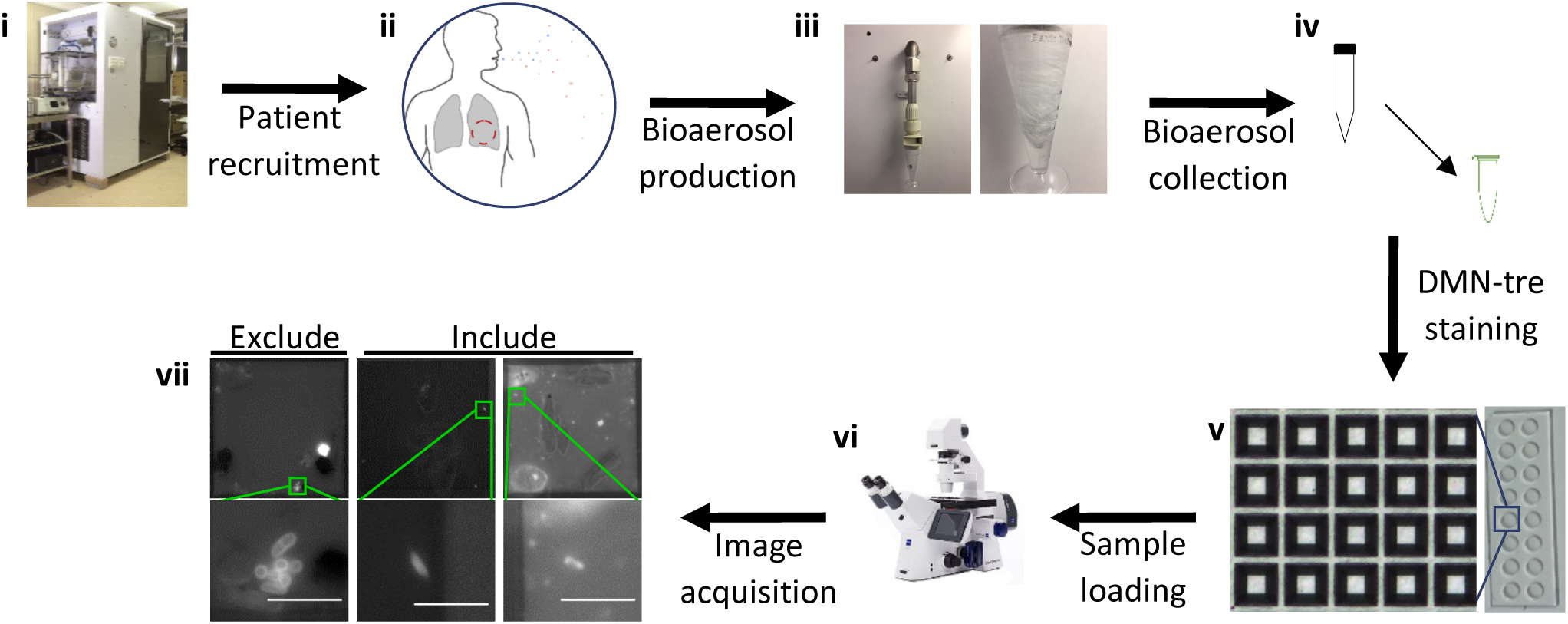
Workflow from participant recruitment to image analysis. (**i**) The Respiratory Aerosol Sampling Chamber (RASC). (**ii**) Bioaerosol production during tidal breathing and non-induced cough. (**iii**) Bioaerosol collection via Bertin Coriolis™ μ Biological Air Sampler. (**iv**) Sample concentration and staining with 100μM DMN-tre during overnight (∼16 hours) incubation at 37 °C. (**v**) The sample is arrayed in the nanowell device. (**vi**) Samples are scanned microscopically and bacilli counted. (**vii**) Relevant data are extracted from images. Scale bar, 5 μm.

The bioaerosol samples of all participants contained large amounts of particulate matter (**Fig. 3**). A qualitative assessment indicated the presence of three broad categories of microparticles/debris: large microparticles with a crystalline appearance were the most common and possibly derived from the Tyvek suits that all participants were required to wear during sampling (**Fig. 3A**); small, auto-fluorescent “splotches”, which were much less common (**Fig. 3B**); and, finally, non-fluorescent and non-staining “granular debris” which was also less common than the large crystalline debris and suggested the presence of human cellular material (**Fig. 3C**).

**Fig. 3.**
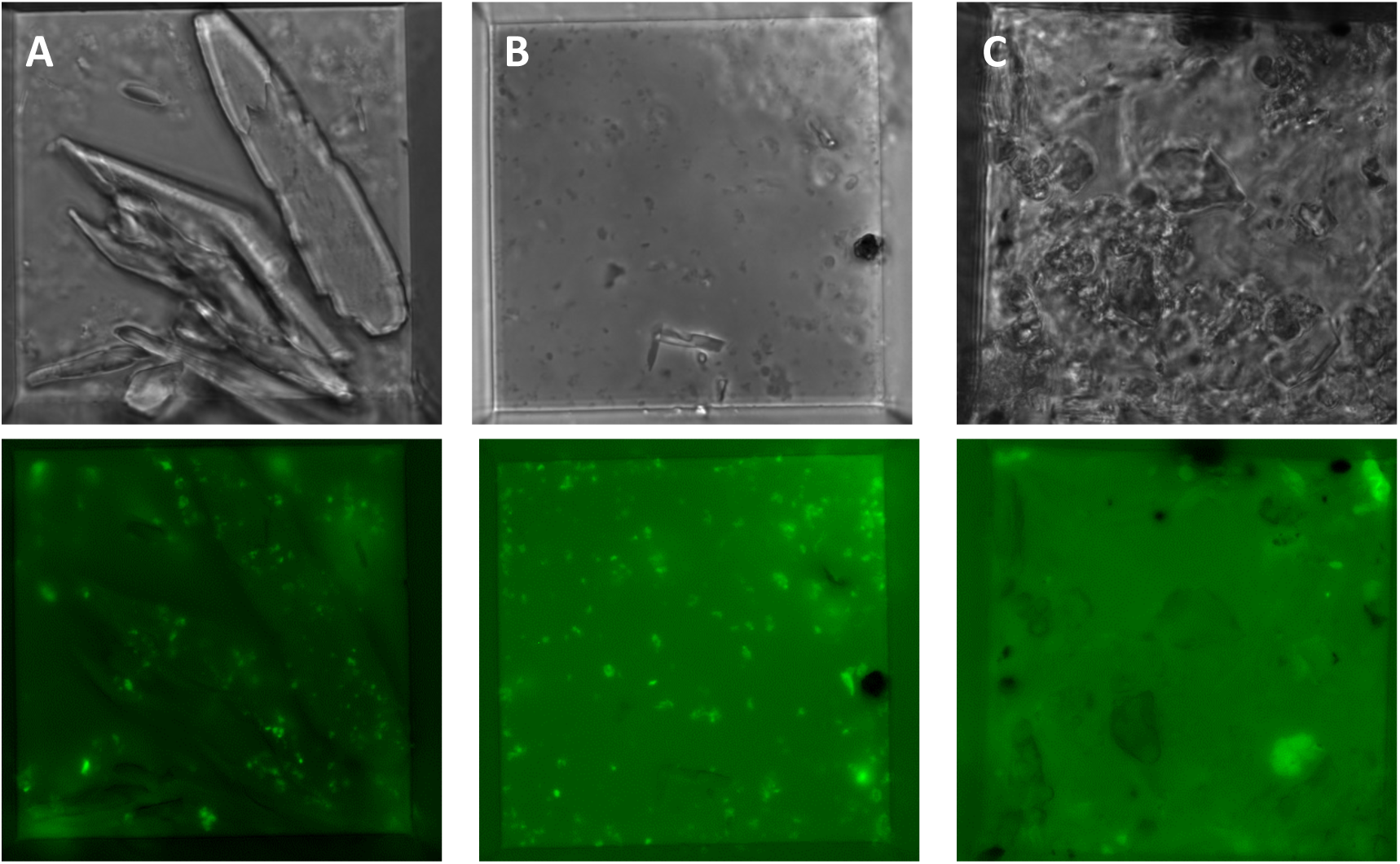
Debris commonly found within TB bioaerosols. Representative images of the three major categories of debris found within bioaerosol samples after overnight staining with 100 μM DMN-tre and visualization within a 50 × 50 μm nanowell. (**A**) Large, crystalline debris, (**B**) small fluorescent debris, and (**C**) granular debris.

The presence of each of these categories of microparticle in any given sample was not mutually exclusive. Every patient produced large micro-particles; however, where the patient samples contained an additional type of debris, it was usually either granular debris or small, fluorescent debris. Further work is required to determine the precise composition and origin of the different micro-particles and, as noted below, to ascertain the relative contributions of auto-fluorescence *versus* true DMN-tre incorporation to some of the fluorescent structures.

### Microscopic investigation revealed the presence of non-Mtb DMN-tre^+^ organisms

The physiological, metabolic and morphological states which characterize *Mtb* during bioaerosol release are not known but are critical to the visual detection of bacilli using the DMN-tre method. To provide a framework (or “identikit”) for the assignment of “putative *Mtb* bacillus” to individual fluorescent structures/bodies detected in the nanowell device, we processed exponentially growing and aged cultures of the laboratory strain, *Mtb* H37Rv, via the clinical DMN-tre labelling algorithm (**Fig. 4A**). The median length of H37Rv bacilli did not change between exponential growth (2.83 μm, IQR = 0.85 μm) and early stationary phase (2.77 μm, IQR = 1.19 μm) (**Fig. 4B**). There was a small but reproducible change in average cell width, with bacilli entering early stationary phase (0.60 μm, IQR = 0.07 μm) very slightly thinner than exponentially growing organisms (0.63, IQR = 0.04 μm) (**Fig. 4B**). However, the average width for these two growth states overlapped substantially (**Fig. 4B**), leading us to reason that any organism falling outside this range (∼0.47 μm to ∼0.86 μm) should be disqualified from classification as “putative *Mtb*”.

**Fig. 4.**
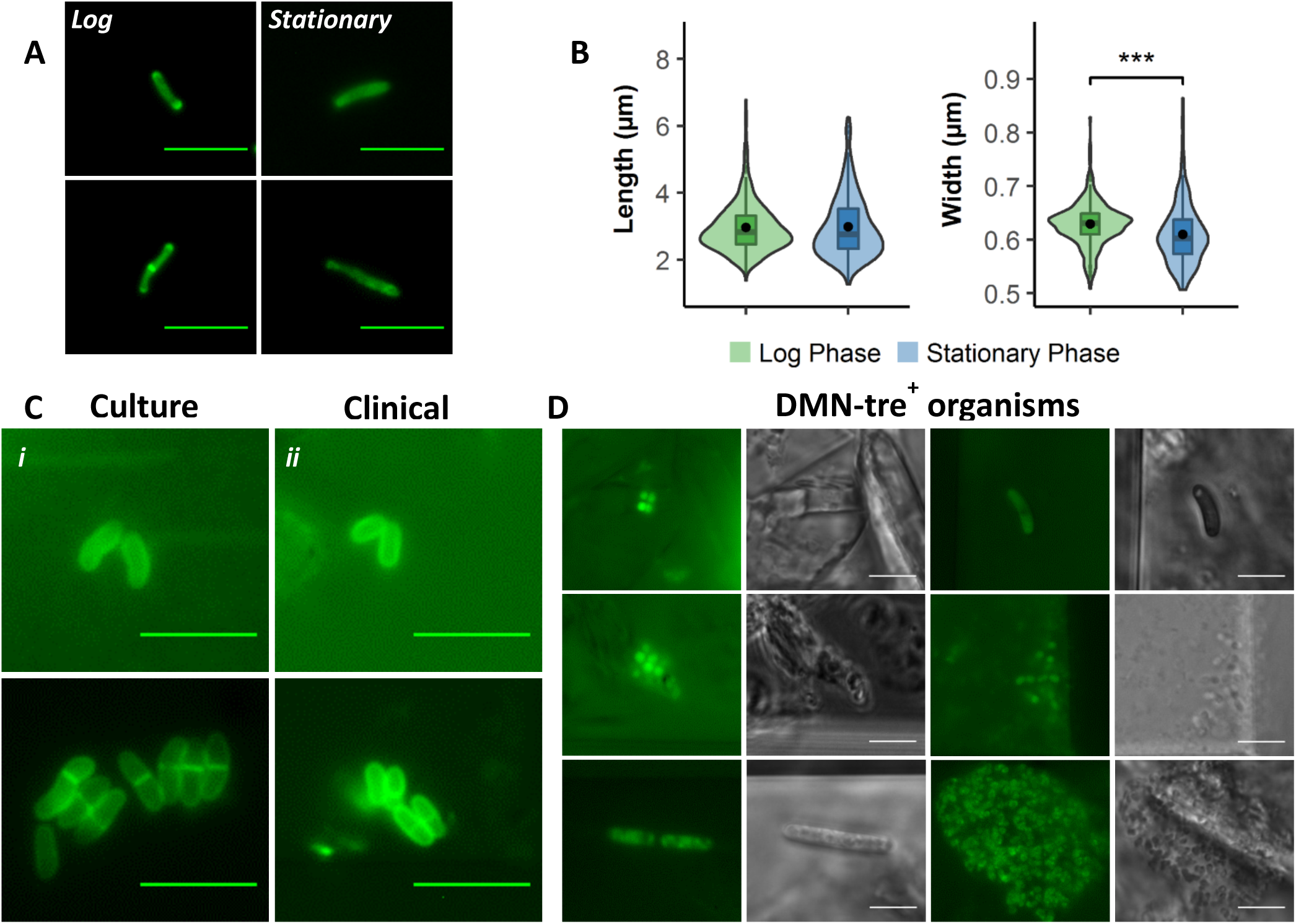
Non-mycobacterial DMN-tre^+^ organisms excluded owing to morphology and staining profile. (**A**) Representative images of *Mtb* H37Rv in log and stationary phase stained with DMN-tre (100 μM) for 2 hrs. (**B**) Plot comparing the length and width of *Mtb* in log-phase (green, n=347) and stationary phase (blue, n = 273). (**C**) Representative images of (i) *C. striatum* cultured in LB broth during log-phase and stained with 100 uM DMN-tre for 5 min, and (ii) DMN-tre positive organisms detected within a bioaerosol sample. (**D**) Corresponding fluorescent and phase contrast images of other non-*Mtb*, DMN-tre^+^ organisms. Scale bar, 5 μm. Wilcoxon Rank-Sum test performed, p < 0.001 (***).

DMN-tre is not specific for *Mtb*: all bacteria within the Actinomycetales encoding homologs of the mycobacterial antigen 85 complex, which catalyzes trehalose mycolylation, possess the capacity to incorporate the fluorescent label. Among these, Corynebacteria are a common constituent of the oral microbiome and have previously been identified in TB sputum samples (*39*). To extend our database of potential DMN-tre-positive organisms, we incorporated the opportunistic pathogen, *C. striatum* (*44*–*46*), in the *in vitro* analyses. Following DMN-tre labelling via the established protocol, *C. striatum* was readily distinguishable from mycobacteria, yielding a distinct cytological (fluorescence incorporation) profile (**Fig. 4C**). Reassuringly, organisms with closely matching features were identified in clinical samples (**Fig. 4C**). However, we also observed multiple DMN-tre^+^ organisms which might be of bacterial and/or fungal origin. Utilizing the exclusion criteria developed above, all of these organisms were eliminated from “putative *Mtb*” classification despite DMN-tre positivity (**Fig. 4C & D**).

### Microscopic identification and characterization of DMN-tre^+^ putative Mtb

Applying the phenotypic classification system, we were able to identify putative *Mtb* in 89 % (25/28) of confirmed TB patients (**Table 1**). Our “empty booth” controls – expected to be *Mtb* free – returned 61 % (14/23) positivity; however, the median count in the TB-positive patients was 15 (range 2-36), whereas the median count in the “positive” empty booths was 3 (range 1-10), suggesting the likelihood that the organisms detected represented carry-over from the cyclone collection system or RASC (**Fig. 5A**).

**Table 1.** Summary of patient characteristics.

|  |  | n (%) | Median count<sup>†</sup> |
| --- | --- | --- | --- |
| Sample type | TB<sup>+</sup> patient | 28 (54.9) | 15.0 |
|  | Empty RASC | 23 (45.1) | 3.0 |
|  | Total | 51 (100) |  |
| Gender | Male | 6 (21.4) | 10.0 |
|  | Female | 7 (25.0) | 7.0 |
|  | NA | 15 (53.6) | 19.0 |
|  | Total | 28 (100) |  |
| HIV status | Positive | 4 (14.3) | 8.0 |
|  | Negative | 9 (32.1) | 7.0 |
|  | NA | 15 (53.6) | 19.0 |
|  | Total | 28 (100) |  |
| Smoking Status | Yes | 7 (25.0) | 7.0 |
|  | No | 5 (17.9) | 7.0 |
|  | NA | 16 (57.1) | 19.0 |
|  | Total | 28 (100) |  |
| Age | Median<sup>††</sup> | 32.5 |  |
|  | IQR | 14.25 |  |

<sup>†</sup>Median number of putative *Mtb* bacilli detected in the sample
<sup>††</sup>15 individuals did not report their age

**Fig. 5.**
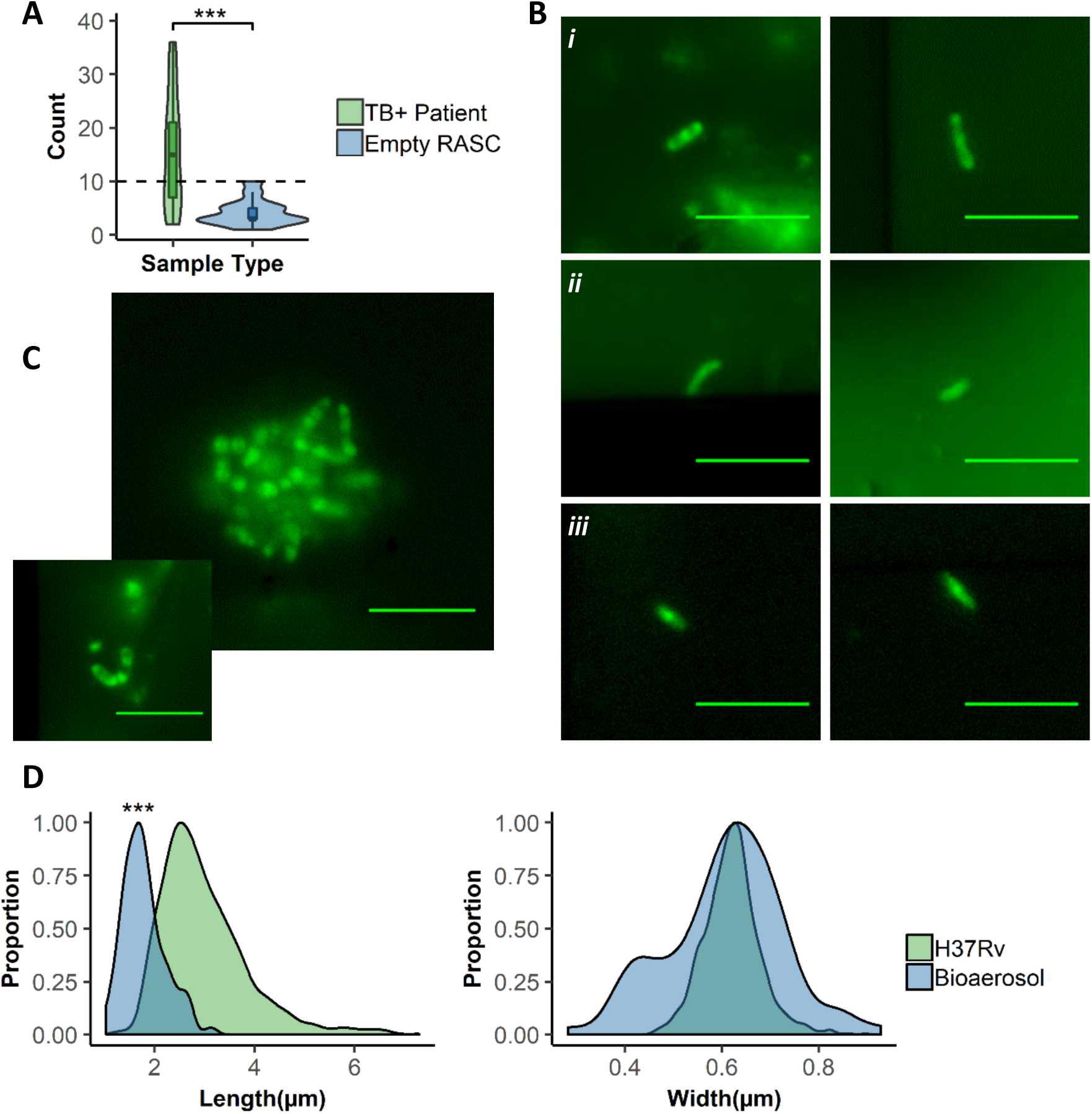
Detection and characterization of putative *Mtb* within bioaerosols of confirmed TB patients. (**A**) Plot comparing the number of putative *Mtb* detected within TB^+^ participants (green, n = 25/28 (89.3 %)) and empty RASC controls (blue, n = 14/23 (60.9 %)). (**B**) Representative images of the three distinct cytological profiles (i-iii) observed in the 25 participants in which putative *Mtb* were detected. (**C**) Representative images of clumps of putative *Mtb* detected within bioaerosol samples. (**D**) Comparing distributions of cell lengths and widths in putative *Mtb* bacilli detected within bioaerosols of TB patients (blue) to *Mtb* H37Rv cultured within the lab (green). Scale bar, 5 μm. Wilcoxon Rank-Sum test performed, p < 0.001 (***).

Three distinct labelling patterns were observed in the putative *Mtb* bacilli that were qualitatively comparable to incorporation patterns previously observed under various culture conditions (**Fig. 5B**). These were: a polar labelling pattern (**Fig. 5B, panel i**), a diffuse labelling pattern (**Fig. 5B, panel ii**), and a labelling pattern which resembled that seen previously in PBS-starved *Mtb* H37Rv *in vitro* (**Fig. 5B, panel iii**). It was notable that polar labelling of bacilli was only observed in patients producing small (auto)fluorescent debris, however the significance (and generalizability) of this observation remains to be determined. In addition, clumps and small clusters of organisms were detected in some samples (**Fig. 5C**). Further characterization of the bioaerosol was possible with single cell resolution. For example, the median cell length was compared for all participant samples in which *Mtb* was detected. These bacilli were on average shorter than in log-phase culture (**Fig. 5D**); in addition, the median length of bacilli found using the liquid sampling method described here was slightly shorter than that observed previously using the Andersen Impactor (*29*).

### Comparison of RASC and sputum profiles

Sputum smear positivity has been a mainstay indicator of patient infectiousness (*47,48*). Moreover, the accessibility and tractability of sputum has enabled numerous investigations of the physiology of infecting bacilli, the key assumption being that the sputum population provides a direct proxy of intrapulmonary and/or transmitted (aerosolized) *Mtb* organisms. To test this notion, we determined whether our sample concentration and DMN-tre labelling process could be applied in parallel to sputum and aerosol populations from the same patient.

An analysis of corresponding sputum and bioaerosol samples from 4 patients suggested a correlation in bacillary numbers between the two samples; however, the small sample size precluded assignment of statistical significance (r = 0.94, p = 0.056) (**Fig. 6A**). Notably, closer inspection of a single patient revealed that each sample was characterized by organisms displaying distinct labelling profiles (**Fig. 6B**). In addition, bacilli within the sputum sample tended to be shorter on average (1.46 μm, SD = 0.585 μm) than those found within the bioaerosol sample (2.21 μm, SD = 0.378 μm) (**Fig. 6B**). Although limited to a single individual, this difference in length might derive from the collection method; that is, it could reflect the fact that aerosolized *Mtb* bacilli might experience a higher level of applied stress (*e.g.* torsional stress) during sample concentration via centrifugation, perhaps resulting in slight filamentation. Alternatively, the differences in staining and cell length might indicate that aerosolized *Mtb* bacilli do not necessarily originate from the sputum, but rather from the peripheral lung, a possibility reinforced by the recent observation that sputum bacillary load may not correlate with the size of the bacillary population captured in face-mask devices (*33*).

**Fig. 6.**
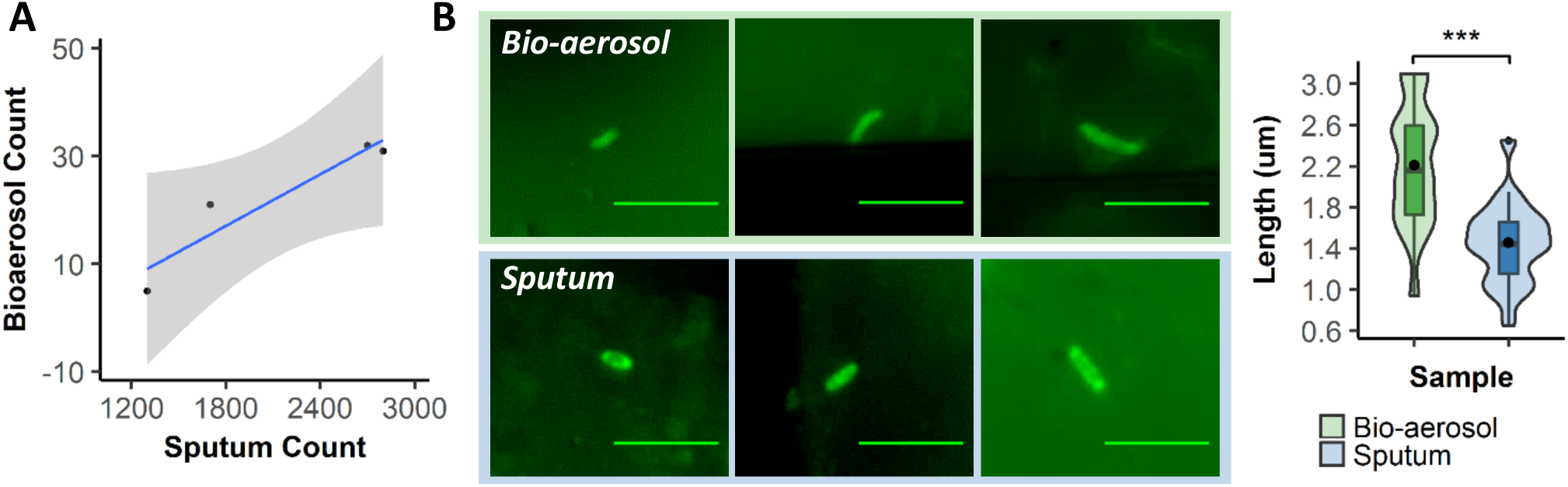
DMN-tre staining and visualisation on the nanowell arrayed slides allows for direct, intra-patient comparison of bioaerosol and sputum samples. (**A**) An intra-patient comparison (n = 4) of the number of putative *Mtb* identified within the bioaerosol compared to sputum (r = 0.94, p = 0.056). (**B**) Representative images from a single patient’s bioaerosol and sputum samples with a comparison of the length between putative *Mtb* bacilli isolated from bioaerosol (green, n = 9) and sputum (blue, n = 38) samples. Scale bar, 5 μm. Wilcoxon Rank-Sum test performed, p < 0.001 (***).

## Discussion

The investigation of TB transmission spans approximately 140 years (*49*). However, until recently, technological limitations had mostly limited microscopic analyses to investigations of the sizes of droplet aerosols. Few attempts have been made to elucidate the physiological and metabolic states of aerosolized *Mtb* bacilli and how these might impact successful transmission to a new host. Similarly, the potential for other material in the aerosol – for example, host-derived cellular and molecular debris, and/or other “contaminating” microorganisms (the “transmission microbiome”) – to influence transmission success is unknown.

Our pipeline for visualization of bioaerosol samples derived from confirmed TB patients has allowed capturing and detection of live putative *Mtb* bacilli in 89% of GeneXpert-confirmed TB patients. From our microscopic analysis, it is evident that the composition of TB bioaerosols is complex and heterogeneous. In designing and implementing the RASC platform, considerable effort was made to minimize the collection of debris (for example, working in a clean-room and requiring that all patients wear Tyvek suits during the sampling process to reduce the production of non-aerosol-derived particulate matter), especially since the cleanliness of samples is critical for reliable microscopic visualization: auto-fluorescence represents a major confounder in environmental samples and there is also the possibility that debris might obscure *Mtb* bacilli partially or even fully. It was for this reason that the nanowell device was utilized, the intention being to maximise *Mtb* detection by increasing the likelihood of separating debris and bacilli through sample dispersal across thousands of nanowell chambers. Of course, it is likely that the composition and origin of the particulate matter might in future prove critical in determining precisely the anatomical origin of aerosolized bacilli. For now, though, raising the signal from putative *Mtb* above the background noise is our priority.

The chemical properties of DMN-tre, most notably the amplification of the fluorescence signal consequent on incorporation of the label into the mycomembrane, ensures its utility in detecting *Mtb* within decontaminated sputum (*39*). However, the fact that Ag85-dependent incorporation of trehalose into the mycomembrane is not unique to mycobacteria (*39*) complicates the application of this label on its own to sign confidently a classification of “*Mtb* positive” bacilli to aerosol samples. At the time this work was initiated, there was a lack of ancillary *Mycobacterium-*specific probes which could be applied to increase the specificity of the microscopic detection while retaining sample viability (thus excluding standard approaches including auramine and acid-fast staining which involve inactivation steps). More recently, other reagents have become available and are under investigation in the RASC platform, including (i) the CDG-DNB3 dual fluorescence probe which is activated by the mycobacterial BlaC β-lactamase and retained in the cell wall following covalent modification by the decaprenylphosphoryl-β-D-ribose 2′-epimerase, DprE1 (*50*); (ii) the TAMRA-labelled benziothiazinone analogue, JN108, which also targets the mycobacterial membrane protein, DprE1, and according to its developers, is able to differentiate *Corynebacterium* from *Mycobacterium* on the basis of fluorescence localization (*51*); and (iii) the Quencher-Trehalose-Fluorophore (QTF) which, like DMN-tre, is a fluorogenic analogue of the natural substrate of mycobacterial mycolyltransferases (*52*). It is too soon to ascertain whether any of these will add value to the analysis; moreover, the overlapping auto-fluorescent signal in many samples urges the development of alternative probes in the red end of the visible spectrum.

The paucibacillary nature of the aerosol samples imposes additional complications which are not applicable to sputum. For example, the processing of bioaerosol samples does not include a decontamination step (sample loss is too significant a factor when dealing with tens of organisms) and, consequently, it is likely that proportionally larger numbers of non-*Mtb* DMN-tre-positive organisms are present in the bioaerosol samples than in decontaminated sputum. For this reason, ongoing work aims to refine our laboratory-based framework for identification of *Mtb* based on metabolic and morphological characteristics. Despite these limitations, we were able to identify with some confidence live putative *Mtb* in 89 % of confirmed TB patients in our study. The detection of putative *Mtb* in “empty booth” controls was most likely due to the carry-over from the cyclone collection system or RASC. Current improvements to the collection system aim to facilitate complete sterilization and elimination of carry-over.

Research over the last 140 years has focused on symptomatic TB diseased individuals as the sole source of infectious particles (*49*). Every year, however, there are more than 10 million new TB cases globally with ongoing *Mtb* transmission the primary driver of incident disease. Despite significant advances in TB diagnostics, immunology and genomic epidemiology (*53*), much remains unknown about individual- and population-level transmission dynamics. This is because our current tools allow for the study of *Mtb* transmission only *after* a TB case is diagnosed. “Community-based” exposure studies focus predominantly on household contacts, which account for less than 20 % of infections in high TB-burden settings (*54–56*). Moreover, the prolonged infectious period of TB (*57–59*) and the potential for transmission from brief, casual exposures mean that fewer than one-third of active TB cases can be epidemiologically and genetically linked (*17,19,20,46,54,60*). In current work, we are adapting the RASC platform to identify viable aerosol production in subclinical cases (*61*), a key departure from prior research which has targeted *Mtb* detection almost exclusively to smear-positive TB.

## Materials and Methods

### Ethics and Patient recruitment

Ethical approval for this study was obtained from the Human Research Ethics Committee at the University of Cape Town (HREC 529/2019). Patients were recruited from two primary healthcare facilities in Masiphumelele and Ocean View, peri-urban townships located very close to each other outside Cape Town, South Africa. Informed consent was obtained from all participants and criteria for inclusion were (i) being 18 years or older, (ii) confirmed GeneXpert positive TB disease, and (iii) no evidence of drug resistant TB. All participants were recruited prior to initiation of standard anti-TB chemotherapy which commenced on the day of diagnosis after sampling was completed.

### Sample collection

Participants recruited into this study were asked to provide two samples: a bioaerosol sample which was collected in the RASC (*37*) and a sputum sample. Bioaerosol collection was done according to previously described protocols (*29*). The sampling method was improved by using a Coriolis μ Biological Air Sampler (Bertin Technologies SAS, France) with the capacity to capture up to 500 L of expired air. Sputum samples were collected in the sputum booth at the clinic.

### Bacterial culture conditions and DMN-tre staining

Culturing of all *Mtb* isolates was conducted in a Biosafety Level 3 (BSL3) facility. *Mtb* H37Rv was grown in Middlebrook 7H9 (Difco™) liquid broth supplemented with 0.2 % (v/v) glycerol, 10% (v/v) Middlebrook OADC enrichment and 0.05 % (w/v) Tween80 (Sigma-Aldrich) at 37°C. *Corynebacterium striatum* was cultured in LB broth (Sigma-Aldrich) at 37°C. For PBS starvation an H37Rv culture was grown to log-phase after which a 1:100 dilution was prepared in fresh 7H9. The diluted cells were harvested by centrifugation (3000 × g for 10 min) and washed three times with an equal amount to original volume PBS + 0.05 % tyloxapol. Following the final wash, the OD_600_ was adjusted to ∼1 and incubated at 37°C for 14 days. For staining of exponentially replicating and stationary-phase bacilli, *Mtb* H37Rv was grown to an OD_600_∼0.5 and ∼1.2, respectively. Bacteria were stained with 100 µM DMN-trehalose for 2h. Stained cells were harvested by centrifugation at 13 000 × *g* for 5 min before resuspending in PBS prior to visualization.

### Staining of samples for visualization in the nanowell device

Staining of all clinical samples was conducted in a Biosafety Level 3 (BSL3) facility.

#### 1) Bioaerosol samples

The 5-10 mL bioaerosol samples were concentrated by spinning at 3000 × *g* for 10 min (Allegra® X-15R, Beckman Coulter). The pellet was resuspended in 200 µl fresh 7H9 and stained overnight (12-16 h), following which the stained sample was concentrated at 13 000 × *g* for 5 min and resuspended in 20 µl sterile, filtered PBS.

#### 2) Sputum

Sputum was treated with 2,3-dihydroxybutane-1,4-dithiol (DTT) (Sigma-Aldrich). An equal volume of 6.5 mM DTT was added to the sputum sample and vortexed until liquified. The liquified sputum was centrifuged at 3000 × *g* for 15 min at 8°C, after which the pellet was resuspended in PBS. To reduce any background host cell material, the MolYsis™ Complete5 kit (Molzym) was used according to manufacturer’s instructions up to the point of bacterial lysis, resuspending the final pellet containing intact bacterial cells in 200 µl 7H9. At this point, the suspension was stained as per bioaerosol staining described above.

Prior to any inoculation, the nanowell device (Edge Embossing) was plasma-coated (Novascan) to counteract the hydrophobicity caused by the polymer used to synthesise the nanowell plate. Samples were loaded into their respective wells and sealed using an adhesive film (ThermoFischer Scientific). The inoculated and sealed plates were centrifuged at 3000 × *g* for 10 min after which they were removed from the BSL3 laboratory for visualization.

### Fluorescence microscopy

Imaging was done on a Zeiss Axio Observer 7 equipped with a 100x plan-apochromatic phase 3 oil immersion objective with a numerical aperture of 1.4. Epifluorescent illumination was provided using a 475 nm LED and non-specific fluorescence was removed, where possible, with a Zeiss 38 HE filter set. Images were acquired using the Zeiss Zen software, and quantitative data were extracted using the MicrobeJ plugin for ImageJ (*62*).

### Statistical analysis

Data were exported from ImageJ and all analysis was performed using R, version 3.5.1. Data normality was assessed visually and, where applicable, a Wilcoxon Rank-Sum test was performed.

## Supplementary Materials

No supplementary materials are provided with this submission.

## Acknowledgments

We are grateful to Lizette Koekemoer for technical assistance in the preparation of starved *M. tuberculosis* H37Rv cultures.

## Author contributions

R.D., S.G., R.W. and D.F.W. designed and led the study. R.W. directed the clinical research site, and C.M. and M.G. assisted with the recruitment of TB patients, ethics application and approval, and transport of the bioaerosol and sputum samples. M.K. and C.R.B provided DMN-trehalose (-tre) reagent and technical input into mycobacterial staining. R.D. performed the BSL2 work and the microscopy. S.G. performed BSL3 experiments, including bioaerosol and sputum sample processing and DMN-tre staining. The nanowell arrayed slides were synthesized by B.S., C.Mc. and A.K., and designed in conjunction with R.D., S.G., and D.F.W. Technical oversite of the RASC was provided by R.W., W.B. and C.C. Data were analyzed by R.D., S.G., and D.F.W. who wrote and edited the manuscript which was approved by all authors..

